# The Breakome of *BRCA1* and *BRCA2* Pathway Mutation Carriers Reveals Early Processes in Breast Oncogenesis

**DOI:** 10.1101/2025.09.25.678591

**Authors:** Sara Oster Flayshman, Osama Hidmi, Jackelyn A. Alva-Ornelas, Jonathan Monin, Mark A. LaBarge, Victoria L. Seewaldt, Yotam Drier, Rami I. Aqeilan

**Author notes:** (R.I.A.), (Y.D.).

## Abstract

DNA double-strand breaks (DSBs) can lead to genomic instability in cancer. Cells rely on an efficient DNA damage response (DDR) to maintain their DNA integrity and prevent oncogenic transformation. However, the early events that connect recurrent DNA damage to oncogenesis are not yet fully understood. Here, using next generation sequencing we comprehensively surveyed genomes to identify DSBs in primary cells of non-malignant carriers of *BRCA1* and *BRCA2* mutations (*BRCA*^mut^), categorized as high-risk patients, to characterize the effects of homologous recombination (HR) loss on cancer initiation. We demonstrate that the landscape of physiological DSBs in *BRCA*^mut^ mammary epithelial cells differs from that of healthy controls and resemble more the DSB pattern observed in breast cancer cells. Our results reveal that proto-oncogenes and tumor suppressors contain more breaks in *BRCA*^mut^ samples, and that genes with a high number of DSBs tend to be more highly expressed. These genes containing a high number of DSBs are also often mutated in breast cancer tumors. Finally, genes with high DSBs in mammary epithelial cells from women with *BRCA*^mut^ exhibit a strong correlation with homologous recombination repair. Together, our findings underscore the impact of *BRCA* loss on the early stages of carcinogenesis and highlight future possibilities for early cancer detection.

**Graphical abstract:** When BRCA is intact, genes that are highly broken are properly repaired via HR, preserving DNA integrity. When BRCA is mutant, impairing its function, highly broken transcriptional DSB genes emerge, no longer able to be efficiently repaired via HR, and are found at genes related to cancer signaling. Breakome of enriched breaks at high-risk model resembles breast cancer breakome, and breaks can be found in genes known to be frequently mutated in breast cancer.

## Introduction

Genomic instability plays a major role in tumorigenesis and is considered a hallmark of biologically aggressive cancers^1,2^. Cells are exposed to a myriad of endogenous (i.e. physiological^3^) and exogenous mechanisms that can result in DNA damage. Factors that can increase DNA damage include 1) replication and transcriptional stress 2) reactive oxygen species (ROS), 3) UV radiation, and 4) other biological and chemical agents^4–6^. The most deleterious form of DNA damage is DNA double-strand breaks (DSBs)^7^; DSBs can lead to severe DNA aberrations and are implicated in cancer. Several DNA damage response pathways are responsible for maintaining genome integrity, specifically homologous recombination (HR) and non-homologous end joining (NHEJ) when DSBs are the culprit^8^. Loss of, or mutations in, DNA repair proteins are known to cause genomic instability and tumor initiation^9–11^.

Genomic instability is tightly associated with breast cancer initiation and progression^12,13^. Approximately 10% of breast cancer is inherited^14,15^; women with deleterious germline *BRCA*1 or *BRCA2* mutations^19–23^ have a lifetime 60-70% risk of breast cancer^14^. *BRCA*1^mut^ carriers most frequently develop highly aggressive triple-negative breast cancer (ER-/PR-HER2-wild type; TNBC; while BRCA2^mut^ carriers most frequently develop luminal (ER+ breast cancers)^16,17^.

Recent efforts have focused on early detection of breast cancers. Next generation sequencing techniques were used to define new cancer genes, mutations, and gene rearrangements^18,19^. DNA damage has been identified through indirect methods, such as staining for DNA-binding damage markers, like γH2AX^20^. While these methods are effective for quantifying relative damage levels, they do not identify specific break sites or the frequencies of specific breaks in the genome. A partial solution is chromatin immunoprecipitation-sequencing (ChIP-seq); however, ChIP-seq remains an indirect method for identifying DNA breaks and lacks high resolution^21^.

Sequencing techniques are being pioneered to directly identify DNA breaks at high resolution^3,22,23^. These approaches show that the landscape of DNA breaks, henceforth referred to as the breakome, is 1) not random and 2) can be used to understand a multitude of DNA breakage and repair processes. Processes that can be better understood include 1) replication mechanisms and 2) high transcriptional stress and resolution. In-suspension Breaks Labeling *in-situ* and Sequencing (sBLISS) is a next generation sequencing (NGS) method that has been developed to detect DNA DSB frequencies at nucleotide resolution ^24–26^. Previous work in our lab using this method helped define the oncogenic role or DNA DSBs in genes that regulate oncogenic transcription, by inducing repair in topoisomerase I regulated transcriptionally active sites, thereby supporting the high burden of oncogenic hypertranscription ^27,28^.

Although the connection between mutations in DNA repair factors and oncogenesis has been long established^29^, how exactly the loss of *BRCA* protein function impacts cancer initiation remains unclear. *BRCA* mutations have been correlated with various known driver events, such as TP53 mutations^30–32^; it is speculated that impaired DNA repair may be responsible for initiating this chain of oncogenic events. Here we investigated how specific *BRCA* mutations impact early changes in DNA break pattern that lead to carcinogenesis. To accomplish this, we collected primary mammary epithelial cells from women with a range of deleterious *BRCA*1 and *BRCA*2 mutations^33^ (Supplementary Table 1). Healthy control samples were obtained from women who are not carriers of deleterious germline DNA mutations, and defined as average risk. Using these samples, we mapped and characterized the pattern of DNA DSBs or “breakome”, and analyzed how specific *BRCA*1 or *BRCA2* mutations may impact early mammary carcinogenesis, and ultimately could play a role in early cancer detection.

## Results

### Normal and *BRCA* mutated primary cells exhibit different break patterns

Given the role of *BRCA* in HR repair, we hypothesized that the break pattern of *BRCA*^mut^ cells would shift to an oncogenic-promoting pattern compared to the normal breast breakome. To understand the effects of *BRCA*^mut^, we subjected our primary cell samples to the sBLISS method ^24–26^ and set out to characterize the breakome of high-risk *BRCA* mutation carriers relative to average-risk controls, without treatment with DNA damage inducers. In total, we analyzed the breakomes of eight heterozygous *BRCA1* high-risk mutation carriers, four BRCA2 heterozygous high-risk mutation carriers, three PALB2 high-risk mutation carriers and seven healthy average-risk controls, all prepared in technical duplicates and were previously confirmed for heterozgosity^33,34^. Tissue samples were collected from several young (≤ 35 years, 4 normal, 5 *BRCA*^mut^), ‘middle aged’ (36–54 years, 2 normal, 5 *BRCA*^mut^) and older (≥ 55 years, 1 normal, 2 *BRCA*^mut^) women of diverse heritages. Samples were labelled with a prefix of either N, B1 or B2, referring to samples that are normal, *BRCA*1 mutated or *BRCA*2 mutated, respectively (Supplementary Table 1). A global view of breaks across the genome shows patterns are largely similar (Supplementary Figure 1A). However, *BRCA* mutated samples tend to be more similar to each other than wildtype, as is evident by Pearson correlation (Supplementary Figure 1B-C) and principal component analysis (Figure 1A). Hierarchical clustering of the whole genome (5677 bins of 0.5MB each) according to Spearman correlation identify two clusters, separating average and high risk almost entirely (Figure 1B). While it is visible that many genomic bins are more broken in high-risk samples, there are also considerable number of bins that are more broken in the average risk samples. Together, we concluded that the breakomes of average-risk and *BRCA*^mut^ high-risk breast epithelia are distinct from one another.

**Fig 1:**
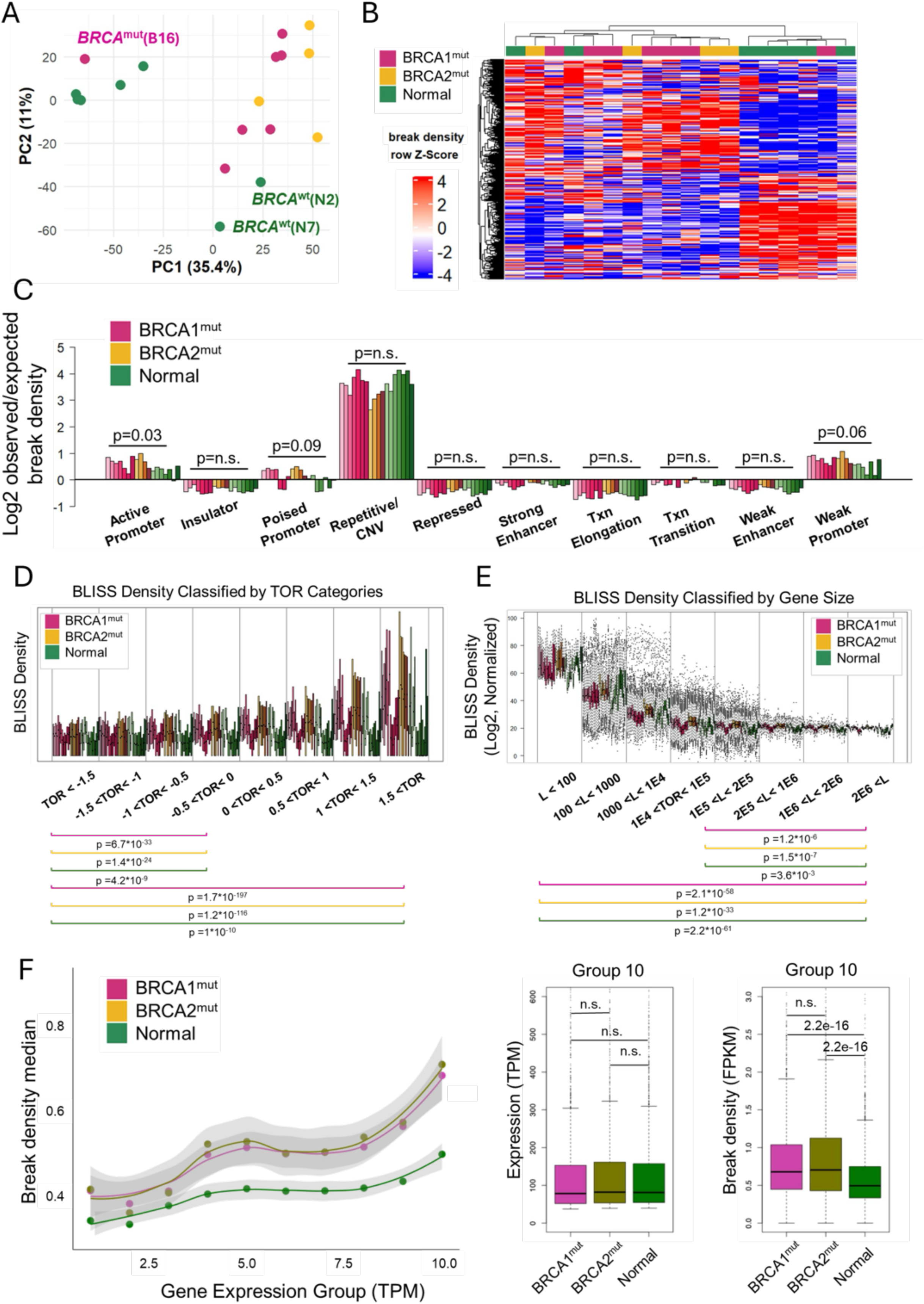
Breakome of *BRCA* Mutated Primary Breast Cells. A. Principal component analysis of the breakome across the genome, binned in 500kbp bins. The analysis demonstrates that *BRCA*-mutated samples exhibit different patterns than average risk samples. B. Heatmap of normalized breaks per 500kbp bin. Rows (bins) and columns (samples) are clustered according to their spearman correlation. Right cluster contains almost only low risk samples, highlighting the unique characteristics of high-risk samples (left cluster). C. Distribution of breaks across chromatin states, bars represent break density in each state and sample compared to the representation of each chromatin state in the genome. P-values were calculated using Kruskal-Wallis similarity test between *BRCA*^mut^ and Normal groups. D. Breaks are enriched in genes that are early replicating. Time-of-Replication analysis depicts gene break density across TOR categories from late replicating genes (left) to early replicating genes (right). P-value was calculated using Kruskal-Wallis similarity test between categories one and four and between categories one and eight for each group (*BRCA1^mut^, BRCA2^mut^, BRCA^WT^*) individually. E. Breaks are enriched in short genes. Gene-length analysis depicts break density across gene length categories from short genes (left) to long genes (right). P-value was calculated using Kruskal-Wallis similarity test between categories one and eight and between categories five and eight for each group (*BRCA1^mut^*, *BRCA2^mut^*, *BRCA^WT^*) individually. F. Breakome vs. expression analysis demonstrates higher break density for genes that are highly expressed. Genes in each expression category were plotted based on their break density levels, each dot representing the median break density per group per expression category. Statistics were measured using Wilcoxon test.

### The breakome of primary cells with *BRCA* mutations is more pronounced in the genome’s open and active regions

To further understand the behavior of physiological DSBs, we analyzed the distribution of breaks in our models across chromatin states as characterized in HMECs. The distribution of observed DSBs relative to the expected bar, which represents the relative distribution of each state across the genome, revealed breaks are overrepresented both at promoters and at repetitive states (Figure 1C). Interestingly the only chromatin state that showed a significant difference between high and average risk samples are active promoters (p<0.03, Kruskal-Wallis similarity test), suggesting breaks in *BRCA*^mut^ are more affected by transcription. Subsequently, we examined the distribution of breaks across genes based on their time of replication. Genes were categorized into groups based on their time of replication, and breaks were counted per gene per sample. We found that early-replicating genes exhibited a higher break density compared to those categorized as late-replicating (Figure 1D). Furthermore, we analyzed the distribution of breaks in genes relative to gene length. Interestingly, shorter genes showed significantly higher break density compared to longer ones (Figure 1E). These findings further suggested that breaks concentrate in highly transcribed regions.

To confirm our hypothesis, we tested whether the breakome correlates with gene expression, utilizing expression data from RNA-seq performed by Shalabi *et al*.^33^ on luminal epithelial cells (LEPs) from the same donors. Genes were binned by their expression levels, starting from 1 for not expressed and going to 10 for highly expressed genes, and break density was measured for genes in each category. Our analysis revealed that higher gene expression correlated with increased break density (Figure 1F). This phenomenon was significantly more prominent in high-risk samples compared to normal breast cells (p < 2*10^-6^, Wilcoxon test), and could also be similarly found when binning genes based on their break density and testing for expression distribution (Supplementary figure 2A, p < 1.7*10^-8^, Wilcoxon test), further highlighting the prominence of transcriptionally mediated breaks in *BRCA* mutated samples.

### DNA DSBs in *BRCA* mutated cells align with cancer pathways from RNA-seq

Next, we shifted to a gene-centric analysis to identify genes more broken in *BRCA*^mut^ samples, potentially due to pathological breaks. We identified 1233 differentially broken genes, comprising 714 more broken and 519 less broken genes (FDR<5%, DESeq2; Figure 2A; Supplementary Tables 2-3). Many of the genes more broken in *BRCA*^mut^ are known proto-oncogenes and tumor suppressor genes (Figure 2B; p < 1.7*10^-6^, one-sided Fisher’s exact test; Supplementary Figure 2B); notably, TP53, an established tumor suppressor, was found to be more broken and is also commonly mutated in *BRCA*1 and *BRCA*2 germline mutated breast cancer^35^. While more broken genes demonstrate higher breakage across the whole gene body, much of the differential breakage is concentrated at the promoter (Figure 2C). To estimate the function of differentially broken genes, we performed gene set enrichment of Molecular Signatures Database (MSigDB). Genes more broken in high-risk samples were enriched for several hallmark gene sets, including *MYC* targets, UV response, *P53* pathway, DNA repair, inflammation, and apoptosis (FDR<25%; Supplementary Table 4; shown FDR<1%, Figure 2D). These pathways not only relate to processes and signaling in cancer, but they also correspond with the pathways enriched in the transcriptome of the same high-risk samples, as previously reported^33^, thereby strengthening our earlier observations regarding the correlation between the breakome and gene expression.

**Fig 2:**
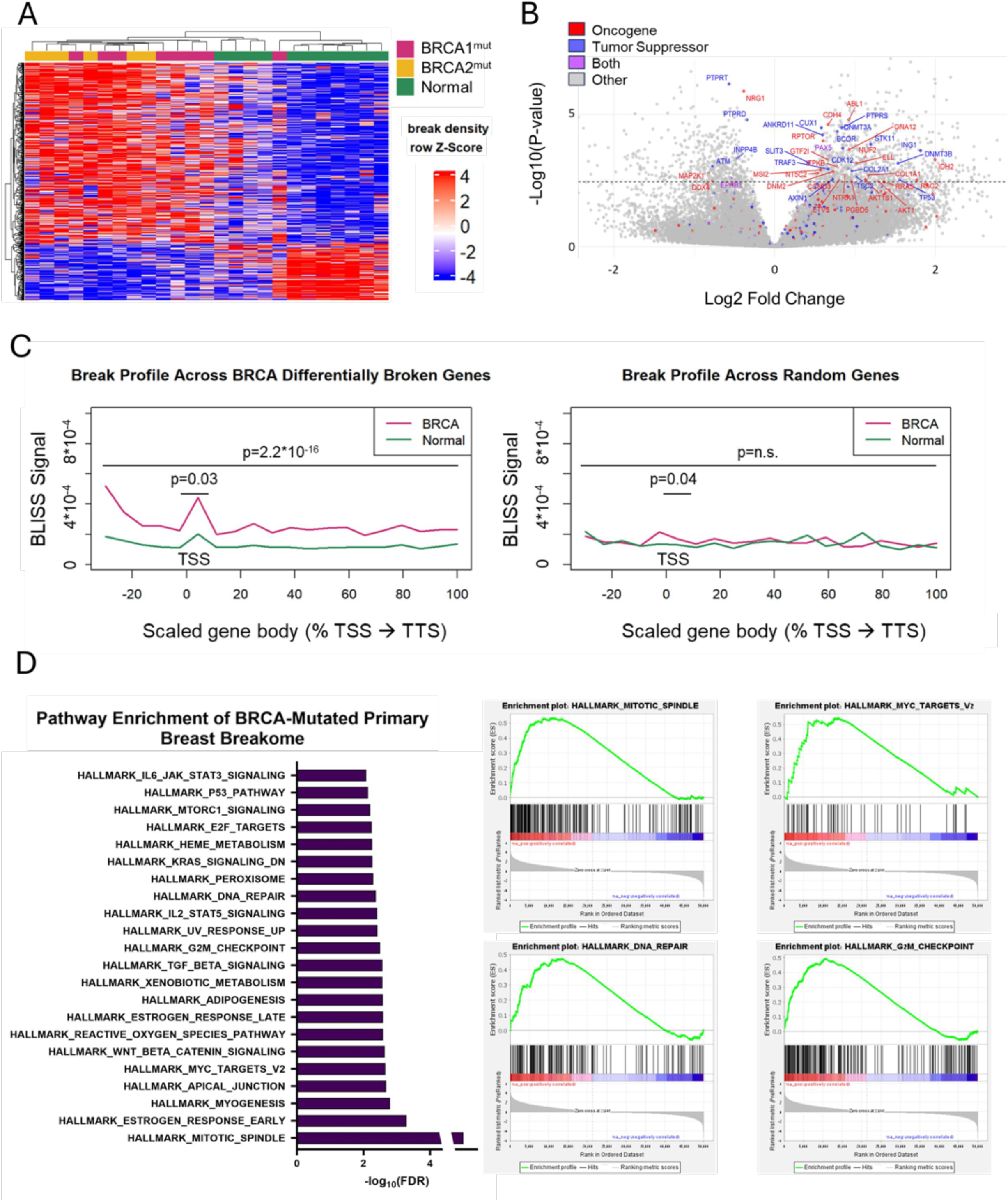
DNA Double Strand Breaks in *BRCA* Mutated Primary Cells Shift Towards Cancer Related Pathways. A. Heatmap of normalized break density at differentially broken genes (FDR < 5%). Rows (bins) and columns (samples) are clustered according to their spearman correlation. B. Volcano plot shows gene break shifts between *BRCA* mutated (positive log2FC) and normal cells (negative log2FC). Dashed line represents threshold of Padj <0.05. Genes were marked and labeled if they met the threshold and considered an Oncogene/Tumor Suppressor in OncoKB database. C. Profile of breaks across gene bodies in differentially broken genes (left) and randomly selected genes (right). Statistics of the region surrounding TSS region were measured using student’s t-test. D. Gene set enrichment analysis of MSigDB pathways enriched in the breakome of *BRCA* samples (FDR < 1%).

### Increased DNA DSBs in BRCA-mutated cells occur at genes undergoing methylation loss

Since methylation status can induce open or closed chromatin, regions of the genome that undergo gain or loss of methylation often overlap with sites exposed to DSBs or other forms of instability, a phenomenon related to shifts in transcription that can act as a catalyst for carcinogenic transformation^36^. To further confirm the relationship between *BRCA* mutated status and transcription, we compared available methylation data overlapping with our gene break data. Indeed, genes which were associated with aging-related methylation loss, and therefore increased expression, according to the data of Senapati *et al*.^37^, demonstrated significantly higher break density compared to genes affiliated with methylation gain, i.e expression downregulation (Supplementary Figure 2C; p < 1.7*10^-6^ for *BRCA*^mut^, p < 0.0005 for *BRCA*^wt^, Wilcoxon test), and especially so for *BRCA*^mut^ samples compared to *BRCA*^wt^ (Supplementary Figure 2C; p < 5.4*10^-16^ for *BRCA*^mut^ vs *BRCA*^wt^ methylation loss, p < 1. 5*10^-13^ for *BRCA*^mut^ vs *BRCA*^wt^ methylation gain, Wilcoxon test). This was also consistent when we tested promoters (Supplementary Figure 2D; p < 0.002 for *BRCA*^mut^, p < 0.0007 for *BRCA*^wt^, p < 1.2*10^-5^ for *BRCA*^mut^ vs *BRCA*^wt^ methylation loss, p < 1.3*10^-5^ for *BRCA*^mut^ vs *BRCA*^wt^ methylation gain, Wilcoxon test). Based on the same data, we also tested the break density of genes specifically associated with transposonal-element related methylation loss, found in DCIS breast cancer. Once again, these genes demonstrated higher break density in *BRCA*^mut^ samples compared to *BRCA*^wt^ (Supplementary Figure 2E; p < 0.02, Wilcoxon test). These findings suggest that *BRCA*^mut^ cells accumulate more DSBs in demethylated, transcriptionally active regions, linking epigenetic deregulation to transcription-associated genome instability.

### DNA DSBs shift between malignant and non-malignant breast cell lines

Previously, sBLISS was utilized to demonstrate various features of cell lines^27,28,38–40^, but a general characterization of breast cell lines is lacking. We therefore applied BLISS to validate malignant and non-malignant breast cancer cell lines (Supplementary Table 5). In line with the literature^27^, the breakome of cell lines was consistent across replicates, while different breast lines exhibited many differences in their breakome (Figure 3A). Principal component analysis further demonstrated this (Figure 3B). Both non-malignant cell lines, HMLE and MCF10a, are similar compared to malignant lines, except for DKAT, a triple-negative breast cancer (TNBC) line^41^ which is characterized by a lack of aneuploidy. MDA-MB231 and MDA-MB436, two additional TNBC lines, showed similar values in their second principal component, as did the two Luminal A lines, MCF7 and T47D. MDA-MB468 is also a TNBC line, yet its breakome resembles Luminal A samples, possibly due to its basal-like morphology, which differs from the mesenchymal-like morphology of MDA-MB231 and MDA-MB436^42^. Hierarchical clustering of the cell lines based on the genome-wide breakome further supports these observations, with replicates clustering together first, non-malignant cells clustering together, and similarity between similar cells such as MDA-MB231 and MDA-MB436 (Figure 3C). The distribution of breaks across chromatin states was consistent with findings in primary cells, demonstrating break enrichment at promoters and repetitive heterochromatic regions, while also being enriched for breaks at strong enhancers (Figure 3D). Furthermore, we tested the correlation of break sites to the H3K4me3 and H3K27me3 histone modifications in each cell line, by creating profile plots in which break loci were centered, and the density of each histone mark was profiled up to 50kb from the break site (Figure 3E). As expected, the enrichment of breaks in promoters is reflected in a strong H3K4me3 peak surrounding break sites. On the other hand, H3K27me3 did not show a consistent pattern around breaks, suggesting polycomb repressed regions are not generally enriched in breaks. Genomic site analysis revealed a high break density for genes in general, particularly in the promoter and 5’UTR regions compared to other genomic features, with intergenic regions being devoid of breaks (Supplementary Figure 3A). As previously observed in primary cells, genes displayed significantly higher break density in early-replicating genes (Supplementary Figure 3B) and in short genes (Supplementary Figure 3C). Together, the characterization of cell lines confirmed our results in primary cells, and allowed us to draw further conclusions using the breakome of breast cancer cell lines as a point of comparison to pre-malignant cell models.

**Fig 3:**
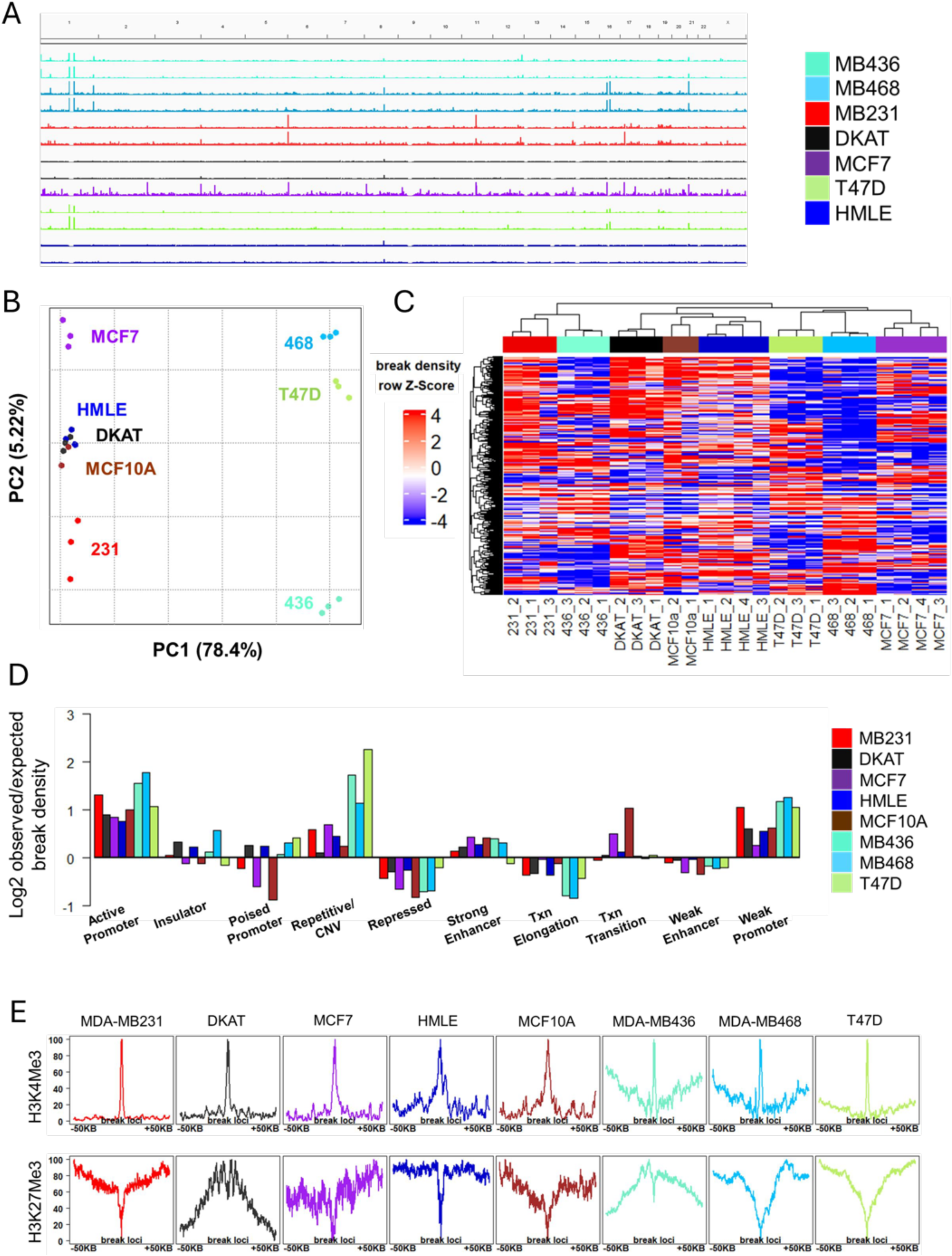
DNA Double Strand Breaks Shift between Malignant and non-Malignant Breast Cell Lines. A. IGV screenshot of genome-wide DSB pattern of breast cell lines, demonstrating similarity within each cell line’s breakome while highlighting differences between distinct cells. B. Principal component analysis of the breakome across the genome, binned in 500kbp bins, of three replicates of each line. C. Heatmap demonstrates sample clustering (spearman) of cell lines based on their binned whole genome. Samples are shown with replicates. D. Distribution of breaks across chromatin states, bars represent break density in each state and sample compared to the representation of each chromatin state in the genome. E. Breaks are enriched at the epigenetic histone mark H3K4Me3, which marks gene expression, and deprived at H3K27Me3, which marks closed chromatin regions. Profile plots encompass the 100 kb region centered by the DSBs, 50kb upstream and 50 kb downstream, while profiling the distribution of the histone mark in that region, relative to breaks.

### High-risk primary cells’ DNA double strand break shifts relate to the breakome of breast cancer cell lines

To test how the breakome of high-risk cells relates to the breakome of breast cancer, we combined our primary cells and cell line data. Primary cells and cell lines display considerable differences, but interestingly, high-risk primary cells were found more similar to the cancer cells than average risk primary cells were (Figure 4A-B). This suggests high-risk samples exhibit an intermediate breakome between average risk and cancer cells.

**Fig 4:**
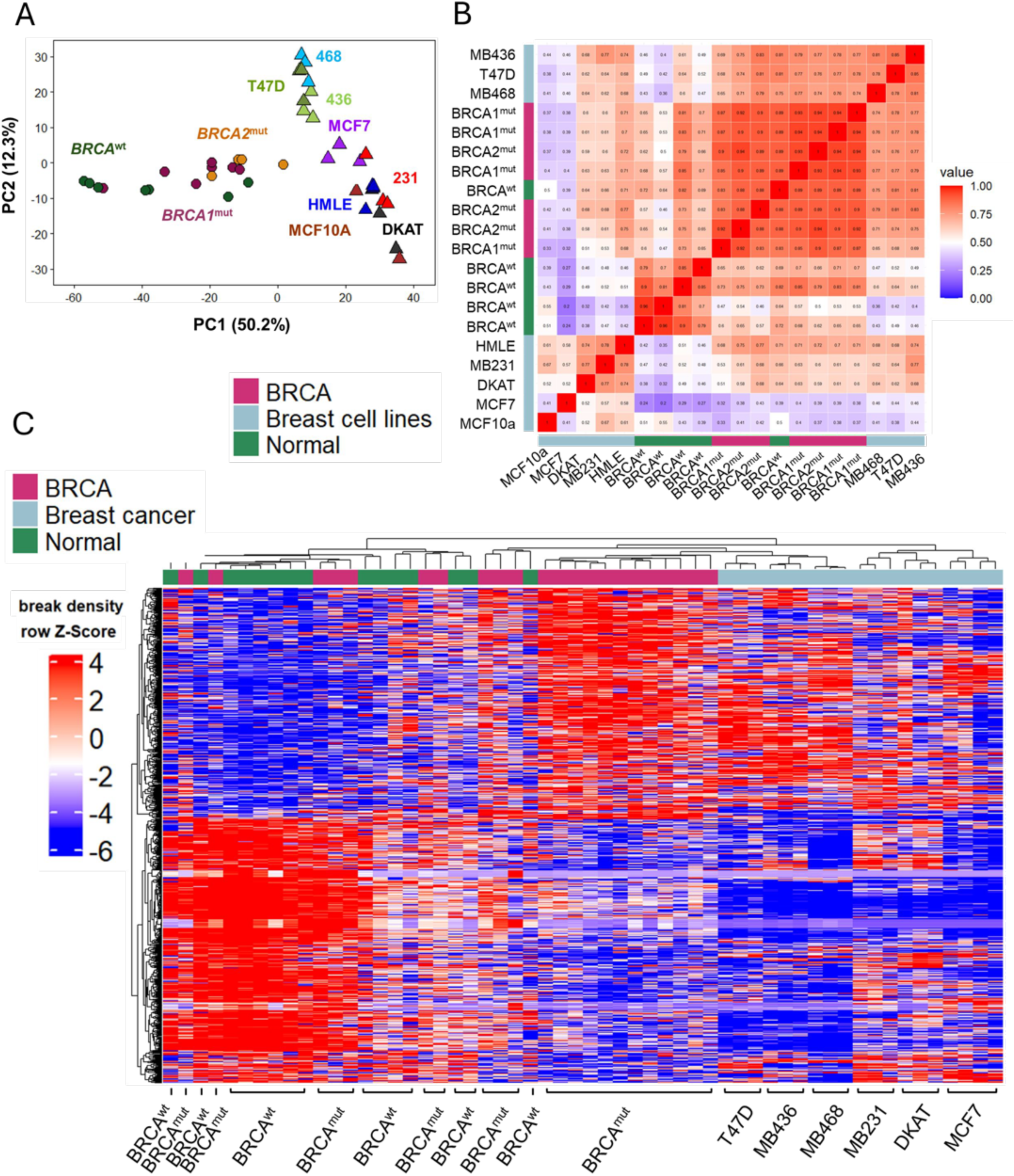
DNA Double Strand Break Shift of High Risk Primary Cells Corresponds to Breast Cancer Cell Line Breakome. A. Principal component analysis of the breakome of normal primary cells, *BRCA*-mutated primary cells and Breast cell lines highlights the intermediate state of high-risk samples, clustering between normal breakome and breast cancer breakome binned whole genome. Samples are shown with replicates. B. Correlation plot highlights similarities and differences between breakome of samples across binned whole genome. C. Heatmap (Spearman hierarchical clustering) of the ‘differentially broken genes’ distinguishing *BRCA* mutated from normal primary cells with inclusion of breast cancer cell line samples demonstrates similar behavior of gene breaks between high-risk primary cells and breast cancer cell lines in genes obtained from DESeq output.

Next, we focused specifically on the list of differentially broken genes between average and high-risk samples. Here, too, malignant cell lines clustered with high-risk *BRCA*^mut^ samples and distanced themselves from normal primary cell samples, demonstrating genes more broken in high-risk samples are also broken in breast cancer cell lines (Figure 4C). Together, these data suggest an intermediary phenotype for high-risk samples, demonstrated by similarities and differences at the level of the breakome, implying certain break patterns emerge very early in carcinogenesis and persist into malignancy.

### The shift in DNA DSBs in high-risk primary cells correlates with mutation frequency in breast cancer and homologous recombination repair

To test if highly broken genes are more likely to accumulate mutations, we utilized breast cancer mutation data from the COSMIC database^43^. Since exonic mutations are subject to selective pressure, we tested mutations appearing in coding sequences separately (Supplementary Figure 4A-C). Consequently, we focused on intronic gene regions, analyzing mutational frequency and focusing on four types of genetic alterations: single-nucleotide variants (SNVs), deletions, insertions and rearrangements. When we analyzed the gene lists for mutational data, overlapping gene break density with gene mutation frequency revealed a higher mutational frequency in genes that were highly broken, for SNVs (Figure 5A), deletions (Figure 5B), insertions (Supplementary Figure 4D) and rearrangements, although not as strongly as the first three (Supplementary Figure 4F). This phenomenon was more prominent and significant in *BRCA*^mut^ cells compared to average risk breast cells (Figure 5A-B and Supplementary Figure 4D,F ; p < 2*10^-4^ for *BRCA*^mut^ SNV, p < 2.35*10^-6^ for *BRCA*^mut^ deletions, p < 3*10^-4^ for average risk deletions, p < 0.04 for *BRCA*^mut^ rearrangements, Wilcoxon test). This enrichment was even more evident when we focused specifically on differentially broken genes. Genes more broken in high-risk cells were enriched in SNVs (p < 2*10^-6^, Wilcoxon test) and deletions (p < 0.019, Wilcoxon test) (Figure 5C) compared to genes more broken in average risk cells. These findings suggest that genes experiencing higher levels of structural disruption in *BRCA*-mutant cells are more prone to accumulating somatic mutations— particularly SNVs and deletions—highlighting a potential mechanistic link between gene fragility and mutational burden in high-risk cancer cells.

**Fig 5:**
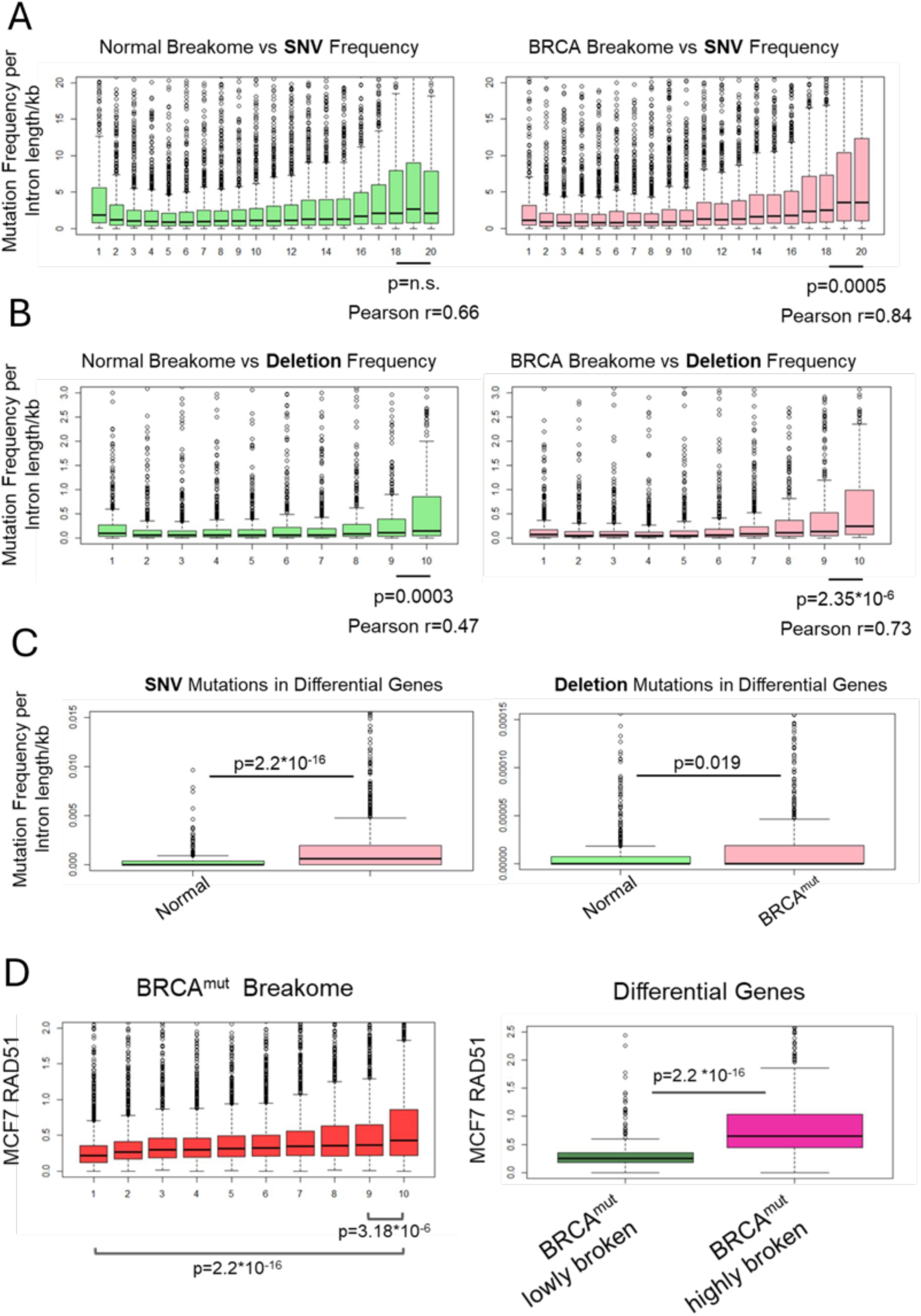
DNA Double Strand Break Shift of High Risk Primary Cells Corresponds to Breast Cancer Mutation Frequency and HR Repair. A. SNV or B. Deletion gene mutation frequency is shown in relationship to normal primary cell gene breakome and *BRCA*-mutated cell gene breakome. Breakome vs. mutation analysis demonstrates a higher mutation frequency for genes that are highly broken. Statistics were measured using Wilcoxon test and Pearson correlation across all of the categories was measured. C. SNV or Deletion gene mutation frequency is higher in *BRCA* mutated Differential genes’ breakome compared to Normal Differential genes’ breakome. Statistics were measured using Wilcoxon test. D. Homologous recombination marker RAD51 enrichment in MCF7 breast cancer associated with gene breakome in *BRCA* mutated samples. Breakome vs. repair analysis demonstrates a higher breast cancer-associated RAD51 binding for genes that are highly broken in *BRCA* mutated cells. Statistics were measured using Wilcoxon test.

To ensure that changes in mutational frequency are not merely a by-product of gene expression levels, we divided the genes into two groups: highly broken/highly expressed and lowly broken/highly expressed. The group with the highest mutational frequency was, in fact, composed of highly broken/highly expressed genes for both SNVs and deletions, while there were no significant differences for insertions between either group (Supplementary Figure 4G). This finding indicates that, although highly broken and highly expressed genes showed the greatest mutagenicity, gene expression alone was not a determinant of breast cancer mutations.

Given the role of *BRCA* in DNA repair, we were curious to explore how repair via HR, or the lack thereof, might contribute to the observed shifts in the breakome landscape of high-risk *BRCA*-mutated breast cells. To investigate this, we utilized ChIP-seq data of the HR DNA repair protein RAD51 in MCF7 breast cancer cells^44^. Interestingly, genes highly broken in *BRCA*^mut^ cells exhibited higher correlation with RAD51 occupancy in MCF7 cells (p < 3.18*10^-6^, Figure 5D). This was even more evident when focusing only on differentially broken genes (p < 2*10^-6^). This suggests these genes may require more HR repair even at a high-risk state, and are therefore more susceptible to mutations, explaining the pattern we observe.

Together, our results demonstrate shifts in the dynamics of DNA repair in cells harboring *BRCA* mutations at very early stages. This might explain why specific genes, shown to be transcriptionally associated, are more susceptible to instability due to the loss of *BRCA* and competent HR. Consequently, these genes, many of which are considered proto-oncogenes and tumor suppressors, are likely to experience persistent breaks and a higher mutation risk, leading those cells down pathways of malignancy.

### DNA DSBs in high-risk *PALB2* mutated cells demonstrate partial resemblance to *BRCA* mutated cells

To understand whether the DSB patterns we observed are unique to *BRCA*1/2 mutations or common to other DNA repair mutations, we performed sBLISS on primary cell samples obtained from three women harboring high-risk *PALB2* germline mutations (Supplementary Table 1). Their break data was compared to specific replicates of an average risk sample (N4), which was prepared and sequenced in the same batch. Similar to what we observed for *BRCA*^mut^, most *PALB2^mut^* samples seem to cluster away from *PALB2^wt^* (Supplementary Figure 5A). Moreover, we searched for differentially broken genes and identified 231 genes more broken in *PALB2*^mut^ samples and 258 less broken genes, but failed to achieve FDR (p < 0.05, DESeq2; Supplementary Figure 5B). While genes that were more broken in *PALB2*^mut^ samples had better overlap with *BRCA*^mut^ samples compared to their WT counterparts, we were not able to identify a significant change in broken tumor suppressors and proto-oncogenes for this sample size (Supplementary Figure 5C-D). Furthermore, we were only able to identify the Hallmark G2M checkpoint pathway as significantly enriched for *PALB2*^mut^ differentially broken genes (GSEA; FDR<25%). Breakome vs. expression analysis recapitulated the notion that genes that are more highly broken demonstrate higher expression distribution (Supplementary Figure 5E). We repeated the analysis on methylation data overlapping with our *PALB2*^mut^ gene break data to confirm whether other HR disrupting mutations would also relate to transcription. Indeed, both for genes and especially promoters, methylation loss genes in *PALB2*^mut^ samples demonstrated significantly higher break density compared to methylation gain genes (Supplementary Figure 5F-G; p < 0.0002 for *PALB2*^mut^ genes, p < 1.2*10^-6^ for *PALB2*^mut^ promoters, Wilcoxon test). *PALB2*^mut^ primary cells and cell lines demonstrated considerable similarities and differences, but not as strongly as we observed for *BRCA*^mut^ (Supplementary Figure 5H). The list of differential genes for *PALB2^mut^* also corresponded with certain breast cancer cell lines (Supplementary Figure 5I). *PALB2*^mut^ break density correlated with RAD51 occupancy in MCF7 cells, as observed previously for *BRCA*^mut^ (p < 1.4*10^-9^, Supplementary Figure 5J). Correspondingly, gene mutation frequency analysis exhibited a higher mutational frequency in more broken *PALB2*^mut^ genes for SNVs, insertions and deletions (Supplementary Figure 5K), which was more prominent in *PALB2*^mut^ than in *PALB2*^wt^ (data not shown). Our data shows that several features of the DSBs seen in *BRCA^mut^* cells also appear in other settings, indicating a broader phenomenon. However, testing these effects in a larger tissue group is necessary to confirm our results.

## Discussion

Women carrying germline *BRCA* mutations exhibit seemingly normal breast morphology^33^, while underlying processes are already in progress. The prevailing dogma is that *BRCA* mutations lead to inefficient DNA repair, resulting in subsequent mutations followed by tumor initiation^45–48^. Despite the roles of *BRCA*1 and *BRCA*2 in HR, and although many have linked deficient repair to mutations, a significant gap in the mechanistic relationship remains unexplored. In this study, we utilized a unique model to capture an understudied time point in carcinogenesis that supports this dogma. For the first time, we demonstrate the distinct biology of non–tumor-bearing, high-risk BRCA mutation carriers and its impact on breast tumorigenesis. While prior studies on related HMEC cohorts have confirmed heterozygosity of BRCA1/2^33,34^, we did not directly assess loss of heterozygosity (LOH) in our samples. Nonetheless, we observed clear shifts in the DNA break pattern compared to non-carrier, average-risk controls. These findings suggest that genomic instability can arise well before LOH, thereby challenging the notion that LOH is the essential initiating event in BRCA-driven tumorigenesis.

Despite that *BRCA*^mut^ samples and average-risk samples cluster almost exclusively away from each other, we still observe a few outliers. Notably, all of the primary cell samples we tested are non-malignant samples, so naturally, despite their important differences, we assume they still have very much in common. We were not yet able to find a factor that would conclusively explain the outlier behavior (age, ethnicity, mutation site or loss of lineage fidelity markers; data not shown), although a larger cohort may be needed to verify any of these hypotheses.

We showed that the breaks resulting from changes in the functionality of *BRCA* are transcriptional DSBs^49,50^, genes that constantly endure exceptional stress from the burdens of the transcription machinery, highlighting their need for effective repair to maintain stability. In fact, we discovered these genes to be enriched for RAD51^45^ in the context of the breast cancer cell line MCF7, with intact *BRCA*.

Many of the genes identified as differential between *BRCA*^mut^ cells and healthy controls are proto-oncogenes and tumor suppressor genes. It is possible to hypothesize that their transcriptional program would shift^46,51^ in response to genomic instability and loss of HR, as a compensatory mechanism to protect the genome, resulting in higher levels of breaks appearing in those regions. This could explain why DNA breaks in normal breast cells do not lead to tumor initiation. Previous work by Shalabi *et al*.^40^ demonstrated a transcriptional shift in *BRCA* mutated cells indicative of an accelerated biological age relative to average-risk women. Exploring the breakome of these older average-risk women might offer further insights into the relationship between transcriptional DSBs and oncogenesis.

Previously, studies aimed to understand cancer mechanisms by applying molecular and OMICS approaches to primary breast cancer samples, cell lines, and manipulated cells^47,48,51–53^. We were interested in determining whether DNA DSBs could explain malignancy by analyzing breakome data of *BRCA*^mut^ in a non-malignant state from authentic carriers. In our work, we were able to show that the shift in broken genes observed in *BRCA*^mut^ cells exhibited a break pattern more similar to breast cancer than to healthy breast cells. This could indicate shifts in transcriptional programs in these cells towards cancer-related pathways^33^. Additionally, detecting breaks that behave similarly to breast cancer at such an early stage led us to hypothesize that DNA breaks might predict the genomic instability landscape typical of breast cancer^53–56^, and suggest likely alterations before they occur. We also showed that higher gene break density in *BRCA*^mut^ cells correlated more strongly with gene mutation frequencies linked with breast cancer, further reinforcing our conclusions regarding the predictive power of the breakome. Higher gene mutation frequencies were detected in most of the mutation types we tested, namely SNVs, deletions, and gene rearrangements. Insertion mutations did not show an advantage in the *BRCA*^mut^ setting when we examined differential genes. This was somewhat surprising, as insertions frequently occur in breast cancer^57,58^. Since insertions alone typically occur in a small number of genes compared to other mutation types, it is possible that the frequencies in our cohort were insufficient to demonstrate any predictive power.

In agreement with our hypothesis, we found that genes which are more highly broken in *BRCA*^mut^ samples also correlate with higher levels of RAD51 occupancy in MCF7 breast cancer cells, which marks repair by HR^45^. More specifically, genes that were differentially broken in *BRCA*^mut^ cells were significantly more enriched for MCF7-RAD51, associating them to repair by HR. Our findings suggest that the absence of *BRCA*1 or *BRCA*2, and thus deficient repair, leads to gene mutation and tumor initiation through the emergence of transcriptionally associated DSBs in genes categorized as tumor suppressors and proto-oncogenes. We also showed that methylation loss–driven transcriptional activation generates fragile genomic regions, and BRCA deficiency exacerbates their breakage due to impaired repair. This reveals an initiation of events occurring at the very early stages of malignancy, highlighting the potential for developing predictive and preventative tools for managing breast cancer diagnosis. Our findings support the notion that in non-cancer carriers, normal but genomically fragile cells may persist in a pre-malignant state, requiring additional genetic or environmental hits to drive their malignant transformation.

An important consideration in this study is the relatively small cohort size (22 samples), which naturally limits the breadth of our conclusions and highlights the need for validation in larger cohorts. In addition, because the tissues were available only as finite HMEC strains, we were unable to perform certain follow-up assays, such as repair protein ChIP-seq, directly on the same samples and therefore complemented our analyses with alternative models. While these comparisons should be interpreted with some caution, the development of future models that permit consistent longitudinal testing would provide an opportunity to further strengthen and extend our findings.

A few questions remain unexplored. For instance, why do germline mutations in *BRCA* only give rise to tumors in specific tissues such as the breast and ovaries, despite affecting DNA repair in all tissues^59,60^. An investigation of the breakome of *BRCA*^mut^ cells in other tissues in relation to transcription could be an interesting step toward further understanding that selectivity. Additionally, pathogenic *BRCA* mutations subject carriers to a 60-80% chance of developing tumors^45,61,62^, raising the question of what is different in the remaining 20-40% of carriers. Many speculate that part of the difference is environmental and hormonal^60,63^, but it is possible that perceptions from the breakome may shed more light on the inherent differences. It would also be interesting and monumental to explore how our findings could translate to the understanding of other cancer types in the future, further expanding prevention and treatment capabilities.

## Methods

### Cell Culture of primary HMECs

Primary HMECs at passage 4 were grown and maintained in M87A medium as previously described^64^. HMECs from reduction mammoplasties were obtained from the HMEC Bank^65^. HMECs from prophylactic mastectomies and tissues contralateral or peripheral to tumors were obtained at City of Hope. Mycoplasma testing was performed on all cell strains before use.

### Cell Culture of cell lines

MCF7 (HTB-22) and T47D (HTB-133) cells were grown in RPMI supplemented with 10% (vol/vol) fetal bovine serum FBS (GIBCO), glutamine, and penicillin/streptomycin. MDA-MB468 (HTB-132) were grown in Leibovitz’s L-15 Medium supplemented with 10% (vol/vol) fetal bovine serum FBS (GIBCO), glutamine, and penicillin/streptomycin. MDA-MB231 (HTB-26) and MDA-MB436 (HTB-130) were grown in DMEM supplemented with 10% (vol/vol) FBS, glutamine, and penicillin/streptomycin. HMLE cells were grown in PromoCell mammary epithelial cell basal media (C-21010) with added supplements (c-93110). MCF10A were grown on DMEM/F12 supplemented with 5% Horse serum, 20 ng/ml EGF, 0.5 mg/ml Hydrocortisone, 100 ng/ml Cholera toxin, 10 mg/ml Insulin and penicillin/streptomycin. Cells were routinely tested for mycoplasma, and cell aliquots from early passages were used.

### In-suspension break labelling *in situ* and sequencing (sBLISS)

sBLISS was conducted as previously described^25,26^. In summary, 10^6^ cells were fixed in 2% paraformaldehyde in 10% FCS/PBS for 10 minutes at room temperature. The fixation was quenched with 125 mM glycine for 5 minutes at room temperature, followed by another 5 minutes on ice and two washes in ice-cold PBS. Cells were lysed for 60 minutes on ice, and their nuclei were permeabilized for 60 minutes at 37°C. Next, nuclei were rinsed twice with CutSmart Buffer containing 0.1% Triton X-100 (CS/TX100), and double-strand break (DSB) ends were blunted in situ using NEB’s Quick Blunting Kit for 60 minutes at room temperature. The blunted nuclei were then washed twice with 1x CS/TX100 before in situ ligation of the sBLISS adapters to the DSB ends. Adaptor ligation was carried out with T4 DNA Ligase for 20-24 hours at 16°C, with BSA and ATP added. Following ligation, the nuclei underwent two washes with 1x CS/TX100, and genomic DNA was extracted using Proteinase K at 55°C for 14-18 hours. Proteinase K was then heat-inactivated for 10 minutes at 95°C, followed by extraction using Phenol:Chloroform:Isoamyl Alcohol, Chloroform, and ethanol precipitation. The purified DNA was sonicated in 100 μL of ultra-pure water using Covaris M220 for 60 seconds. Sonicated samples were concentrated with AMPure XP beads (Beckman Coulter), and fragment sizes were evaluated using a BioAnalyzer 2100 (Agilent Technologies), targeting a range of 300 bp to 800 bp with a peak around 400-600 bp. The sonicated DNA was then in vitro transcribed using the MEGAscript T7 Kit for 14 hours at 37°C. After RNA purification and ligation of the 3’-Illumina adaptors, the RNA underwent reverse transcription. The final library indexing and amplification step was performed with NEBNext® Ultra™ II Q5® Master Mix.

sBLISS fastq files are initially de-multiplexed using sample barcodes. Quality control is performed with trim galore to eliminate residual adapters, trim reads to a base quality of at least 20, and filter out short reads smaller than 20 bp. Initial and final sample qualities are assessed with fastqc. Quality-processed fastq files are aligned to the GRCh38 assembly with hisat2, then sorted and indexed using samtools. The resulting bam files are de-duplicated with umi-tools, utilizing genomic coordinates and Unique Molecular Identifiers (UMI). Custom Python and R scripts are employed to identify read start positions and convert bam files to bigwig format and BED files for downstream analysis. An additional custom R script is used to discard a blacklist of positions, primarily within centromeres.

BED format Breakome data was associated with either genes or 0.5MB bins using Bedtools V2.26.0^66^. For genes, breaks are counted per gene based on overlapping annotations, GRCh38 gene annotations were downloaded from GENCODE^67^. For bins, whole genome is tiled in windows of 0.5MB each and breaks are counted per tile. Data was subsequently normalized for FPKM or TPM and Z-score. In some cases, break data was averaged across all samples of the same group and subsequently FPKM or TPM normalized. Further details can be found in figure legends.

Principal component data was obtained using prcomp function of the R stats base package and visualized using the R ggplot2 package^68^.

Heatmaps were created with the ComplexHeatmap R package^69,70^.

‘Differential break’ analysis was performed using DESeq2^71^ in R v.4.2. Gene set enrichment for MsigDB hallmark pathways was subsequently performed on the DESeq output using GSEA 4.3.2 for pre-ranked data, by their ‘stat’ column. Genes were analyzed for corresponding pathways using the built in chip platform MSigDB v.2023. Ranking method was signal to noise and hits were considered significant if FDR<0.25.

Profile plots were created using BigWig data. Files were imported using R package Rtracklayer^72^, normalized and averaged per group for *BRCA*^mut^ or normal groups. List of genes to profile was subsetted for either positively-differentially broken or random list and then scaled into 10 bins for each gene with a 25% extension before the TSS to capture the full promoter region. Breaks were then profiled for each group across the gene representing region.

### Chromatin states

Chromatin states for HMEC were obtained from the chromatin state segmentation ChromHMM by ENCODE (GSE38163) as described^73,74^. States used for plotting are as follows: 1_Active promoter, 8_Insulator, 3_Inactive/poised Promoter, 14/15_Repetitive/CNV, 12_Polycomb-repressed, 4/5_Strong Enhancer, 10_Transcriptional Elongation, 9_Transcriptional Transition, 6/7_Weak/poised Enhancer, 2_Weak promoter. Normalized breaks were counted per selected chromatin state for each sample. The percentage representation of each chromatin state in the genome was determined, then Log2 observed/expected was calculated.

### Time-of -Replication

Samples were analyzed for TOR data downloaded from ReplicationDomain software^75^. Replication timing used was as previously defined by breast cancer MCF7 cell line. Genes were grouped based on their replication timing category and normalized break density was plotted.

### RNA-seq data

RNA-seq data from primary HMECs was performed at City of Hope and is available at GSE182338. Only samples that matched the ones we possessed were used and the analysis was performed on the LEP data. TPM values were averaged across replicates for each group to determine the expression rank of each gene. Further analysis of the RNA-seq data can be found at *Shalabi et al. Nature Aging, 2021*^33^.

For expression vs. breakome analysis, 22130 TPM normalized genes were divided into 10 categories based on their TPM expression per group (*BRCA1^mut^*, *BRCA2^mut^*, *BRCA^WT^*), category 1 being the least expressed, up to category 10, highly expressed. Genes in each category were plotted based on their break density levels, each dot representing the median break density per group per expression category.

For combined breakome and expression vs. mutation analysis, samples’ breakome data were merged for *BRCA^mut^* and TPM normalized. Genes were considered highly or lowly broken if their TPM break levels belonged to the top or bottom 10% breakome, respectively. For Expression categories, samples were merged for *BRCA^mut^* and TPM normalized. Genes were considered highly expressed if their expression levels were TPM>20. Genes in each of the two categories had to meet both mentioned conditions. Then, mutation frequency was plotted for the genes in each group.

### ChIP-seq data

Modifications H3K4me3 and H3K27me3 were downloaded from ChIP-Atlas^76^ (GSE85158^77^) based on hg38 genome build. Top scored 1000 breaks in each cell line were selected, and histone modification peaks were examined in the +/-50Kbp vicinity of these breaks.

ChIP-seq data for RAD51 in MCF7 cells were downloaded from ChIP-Atlas (GSE105597^44^), and accumulated signal was associated with genes based on their annotations as mentioned above and FPKM normalized. Samples were merged for each group (BRCA^mut^ or normal) and TPM normalized, then split into 10 categories based on their break density, category 1 being the least broken, up to category 10, the most broken. Genes in each category were plotted based on their RAD51 signal. For differential genes analysis, genes were grouped as either ‘Normal’ or ‘*BRCA*^mut^’ based on DESeq output (Normal, log2FC<0 and Padj<0.05; *BRCA*^mut^, log2FC>0 and Padj<0.05). Then, RAD51 signal was plotted for each group.

### SNV, Deletion and Insertion mutations data analysis

The file Cosmic_NonCodingVariants_Tsv_v101_GRCh37.tar, containing all noncoding variants, was downloaded from COSMIC. This file was first filtered for breast cancer samples and separated based on the type of mutation into SNVs, deletions, and insertions mutation files. These files were then converted into genomic ranges objects and filtered to contain only mutations within introns. To calculate mutation density, mutations per gene were counted and normalized by the sum of intron length.

For Breakome vs. mutation analysis, breakome categories were created by merging samples for each group (BRCA^mut^ or normal), TPM normalizing, then splitting genes into 20 or 10 categories based on their break density, category 1 being the least broken, up to category 20/10, the most broken. Per mutation type, mutation frequencies of breast cancer related genes (either coding or intronic regions) were normalized to kb. Genes in each break category were plotted based on their mutation frequencies.

For differential genes analysis, genes were grouped as either ‘Normal’ or ‘*BRCA*^mut^’ based on DESeq output (Normal, log2FC<0 and Padj<0.05; *BRCA*^mut^, log2FC>0 and Padj<0.05). Then, mutation frequency was plotted for each group.

### Rearrangement mutations data analysis

Mutation data was obtained from Nik-Zainal *et al*. 2016^78^. The list of genes and their respective rearrangement mutation densities were taken as is from table 13 in the publication. Breakome data was prepared the same as for the analysis of SNVs, deletions, and insertions. Genes in each break category were plotted based on their mutation density.

### Methylation data

Methylation data of aging primary cells was obtained from Senapati et al^37^. The list of genes and their respective methylation status were taken as is from table 5 (5A for methylation loss status and 5B for methylation gain status) in the publication. The gene list for DCIS age-dependent methylation loss was taken as is from table 9 in the publication.

### Statistics

Groups were compared for normal distribution and variance, to determine relevant statistical test. Figure legends list the statistical test used during the experiment. Unless otherwise specified, all tests reported were two tailed. R v.4.2 software was used for statistical analysis. Significance was achieved when *P* < 0.05 or Binjamini-Hochberg adjusted *P* < 0.05 when testing for multiple hypotheses. In the case of GSEA, significance of an enriched pathway was achieved when FDR < 0.25.

## Acknowledgments

We sincerely thank the members of the Aqeilan lab for their valuable discussions and insights. We are deeply grateful to Prof. Itamar Simon for his thoughtful guidance and support as a member of the PhD committee of S.O.F. We also appreciate the invaluable assistance of Dr. Abed Nasereddin and Dr. Idit Shiff from the Core Research Facility of the Hebrew University-Hadassah Medical School. This study was supported by grants from the Israel Science Foundation (ISF) [No. 1056/21]. We also acknowledge the support of the Carole and Andrew Harper Diversity Scholarship Program and the VATAT PhD scholarship to O.H. and the Science Training Encouraging Peace (STEP) Fellowship previously supporting S.O.F.

## Human subjects statement

Samples were collected under City of Hope IRB #17185. The trial is subject to United States federal regulations concerning clinical trials and undergoes yearly review; the trial is currently active. Subjects were consented in writing and in person.

## Author contributions

S.O.F., Y.D. and R.I.A. conceived and designed the study. S.O.F performed all sBLISS experiments. S.O.F, J.M. and O.H. performed computational analysis. S.O.F, O.H., J.A.A.O, V.E.S. and M.A.L., Y.D. and R.I.A wrote the manuscript. J.A.A.O, V.E.S. and M.A.L. provided the *BRCA*^mut^ and *BRCA*^wt^ primary cell samples. J.A.A.O_grew the primary cell samples and fixated them for sBLISS by S.O.F.

## Data Availability

sBLISS data have been deposited in the Gene Expression Omnibus database under accession no. GSE300163.

---

**Supplementary Fig 1:** The Breakome of Normal and *BRCA* Mutated Primary Cells can be Distinguished A. Integrative genomics viewer (IGV) software was used to visualize the genome-wide DSB pattern of primary breast cells in BigWig format, and demonstrates similarities and differences between the breakome of normal primary breast cells and *BRCA*-mutated ‘high risk’ primary breast cells. B. Correlation plot (Pearson) highlights similarities and differences between breakome of samples across binned whole genome. C. Correlation-based similarity index represents the mean correlation between each group (whithin *BRCA*^mut^, within Normal) and the correlation between the two groups. Correlation values were averaged for each group and plotted as shown. Statistics were calculated using Wilcoxon test.

**Supplementary Fig 2:** Genes more broken in *BRCA* mutated cells are transcriptionally associatedA. Breakome vs. expression analysis demonstrates a higher expression distribution for genes that are highly broken, more prominently so in *BRCA* mutated samples. Genes in each break category were plotted based on their expression levels. Statistics were measured using Wilcoxon test. B. OncoKB tumor suppressors and proto-oncogenes are more susceptible to breakage in *BRCA* mutated samples. Highly or lowly broken were defined based on the differential breakage of *BRCA* mutated cells. Statistics were measured using Wilcoxon test. C. Methylation loss genes and D. Methylation loss promoters demonstrate higher break density in *BRCA* mutated cells compared to average risk controls at a higher level than observed in methylation gain genes/promoters. Statistics were measured using one-tailed Wilcoxon test. E. *BRCA* mutated samples have higher break density in genes which are associated with methylation loss in breast DCIS tumors. Statistics were measured using one-tailed Wilcoxon test.

**Supplementary Fig 3:** Characterization of DNA double strand breaks across breast cell lines A. Distribution of breaks across genomic sites shows break enrichment across genes and especially at the 5’ UTR and promoter regions. Bars represent the normalized break count enrichment per selected site for each cell line. ‘Expected’ depicts the representation of each site in the genes across the genome, making it the point for comparison. B. Breaks are enriched in genes that are early replicating. Time-of-Replication analysis depicts break density across TOR categories from late replicating genes (left) to early replicating genes (right). P-value was calculated using Kruskal-Wallis similarity test. C. Breaks are enriched in short genes. Gene-length analysis depicts break density across gene length categories from short genes (left) to long genes (right). P-value was calculated using Kruskal-Wallis similarity test.

**Supplementary Fig 4:** DNA Double Strand Break Shift of High Risk Primary Cells Corresponds to Breast Cancer Mutation Frequency and HR Repair A. SNV, B. Deletion or C. Insertion coding gene mutation frequency is shown for differential genes. SNV and Deletion coding gene mutation frequency is higher in *BRCA* mutated Differential genes’ breakome compared to Normal Differential genes’ breakome. Statistics were measured using Wilcoxon test. D. Insertion gene mutation frequency is shown in relationship to normal primary cell gene breakome and *BRCA*-mutated cell gene breakome. Breakome vs. mutation analysis demonstrates a higher mutation frequency for genes that are highly broken in BRCA^mut^ samples. Statistics were measured using Wilcoxon test and Pearson correlation across all of the categories was measured. E. Insertion gene mutation frequency shows no significant advantage in *BRCA* mutated Differential genes’ breakome compared to Normal Differential genes’ breakome. Statistics were measured using Wilcoxon test. F. (same as D) Normal and *BRCA*-mutated Breakome vs. mutation analysis was similarly preformed for rearrangement mutation density. Statistics were measured using Wilcoxon test and Pearson correlation across all of the categories was measured. G. Gene mutation frequency is shown relative to BRCA^mut^ Breakome and Expression categories (Breakome^high^Expression^high^ or Breakome^low^Expression^high^), to show that expression alone is not a determiner of mutation. Statistics were measured using Wilcoxon test.

**Supplementary Fig 5:** Samples with germline *PALB2* mutations considered high risk demonstrate a partially similar phenotype to *BRCA* mutated samples. A. Principal component analysis of the breakome across the genome, binned in 500kbp bins. The analysis demonstrates that most *PALB2*-mutated samples exhibit different patterns than average risk samples. B. Heatmap of normalized break density at differentially broken genes (P < 5%). Rows (bins) and columns (samples) are clustered according to their spearman correlation. C. Venn diagram shows more genes in common between differentially highly broken genes in *BRCA* mutated samples and *PALB2* mutated samples compared to lowly broken genes. D. Volcano plot shows gene break shifts between *PALB2* mutated (positive log2FC) and normal cells (negative log2FC). Horizontal dashed line represents threshold of P <0.05. Vertical dashed line represents Log2 fold change =1. Genes were marked and labeled if they met the threshold and considered an Oncogene/Tumor Suppressor in OncoKB database. E. Breakome vs. expression analysis demonstrates a higher expression distribution for genes that are highly broken in *PALB2* mutated samples. Genes in each break category were plotted based on their expression levels. F. Methylation loss genes and G. Methylation loss promoters demonstrate higher break density in *PALB2* mutated cells compared to average risk controls at a higher level than observed in methylation gain genes/promoters. Statistics were measured using one-tailed Wilcoxon test. H. Correlation plot highlights similarities and differences between breakome of samples across binned whole genome. I. Heatmap (Spearman hierarchical clustering) of the ‘differentially broken genes’ distinguishing *PALB2* mutated from normal primary cells with inclusion of breast cancer cell line samples demonstrates a partially similar behavior of gene breaks between high-risk *PALB2*^mut^ primary cells and breast cancer cell lines in genes obtained from DESeq output. J. Homologous recombination marker RAD51 enrichment in MCF7 breast cancer associated with gene breakome in *PALB2* mutated samples. Breakome vs. repair analysis demonstrates a higher breast cancer-associated RAD51 binding for genes that are highly broken in *PALB2* mutated cells. Statistics were measured using Wilcoxon test. K. SNV, Insertion or Deletion gene mutation frequency is shown in relationship to PALB2-mutated cell gene breakome. Breakome vs. mutation analysis demonstrates a higher mutation frequency for genes that are highly broken. Statistics were measured using Wilcoxon test and Pearson correlation across all of the categories was measured.

